# Genome-guided vs transcriptome-guided mapping: systematic evaluation of RNA-seq quantification bias for tandemly duplicated genes in model and crop plants

**DOI:** 10.64898/2026.11.24.768806

**Authors:** Yuxuan Jing, Fei Wang, Jingwei Ma, Yan Bao, Peipei Wang

## Abstract

Accurate quantification of RNA-seq data is fundamental to modern plant biology. The choice of mapping strategy and aligner critically determines expression quantification accuracy and shapes the quality of downstream biological analyses. However, a pervasive yet often underappreciated challenge is the accurate quantification of genes located within tandemly duplicated arrays. Here, we selected six representative plant species—*Arabidopsis thaliana* and five important crops, and compared RNA-seq analysis results generated from two mapping strategies (genome-guided and transcriptome-guided) using three widely used aligners (TopHat2, HISAT2, and STAR). Our results reveal pervasive alignment errors in genome-guided strategies, arising primarily from misaligned reads spanning tandem gene duplicates—a gene class that plays important roles in plant defense and stress responses, detoxification, and secondary metabolism. TopHat2 with the transcriptome-guided strategy yielded the most accurate alignments, whereas transcriptome-guided HISAT2 and STAR corrected most, but not all, of the errors. Transcriptome-guided mapping also demonstrates superior performance over the genome-guided approach on simulated RNA-seq benchmark datasets. Moreover, in functional enrichment analysis, the transcriptome-guided strategy outperformed the genome-guided one in retrieving gene sets biologically relevant to the processes under study. Overall, this study provides a systematic evaluation of how mapping strategies and software selections affect RNA-seq quantification for tandem gene duplicates, and offers a valuable reference for standardizing plant RNA-seq analytical workflows, particularly for studies investigating biology processes regulated by tandem gene duplicates.

## Introduction

Transcriptomic analysis is critical for advancing biological research, serving as an essential tool for decoding cellular physiological and pathological responses at the molecular level (Wang *et al*., 2009). Since the development of next generation sequencing technologies around 2005, publications involving transcriptomic analysis have increased rapidly (**Fig. 1a**). The accuracy and reliability of this analysis fundamentally depend on the precise alignment of RNA-seq reads to a reference genome or transcriptome. Mapping tools such as TopHat (Trapnell *et al*., 2009)/TopHat2 (Kim *et al*., 2013), Hierarchical Indexing for Spliced Alignment of Transcripts (HISAT) (Kim *et al*., 2015)/HISAT2 (Kim *et al*., 2019), and Spliced Transcripts Alignment to a Reference (STAR) (Dobin *et al*., 2013) have been developed to align the RNA-seq reads (**Fig. 1b**).

**Fig. 1.**
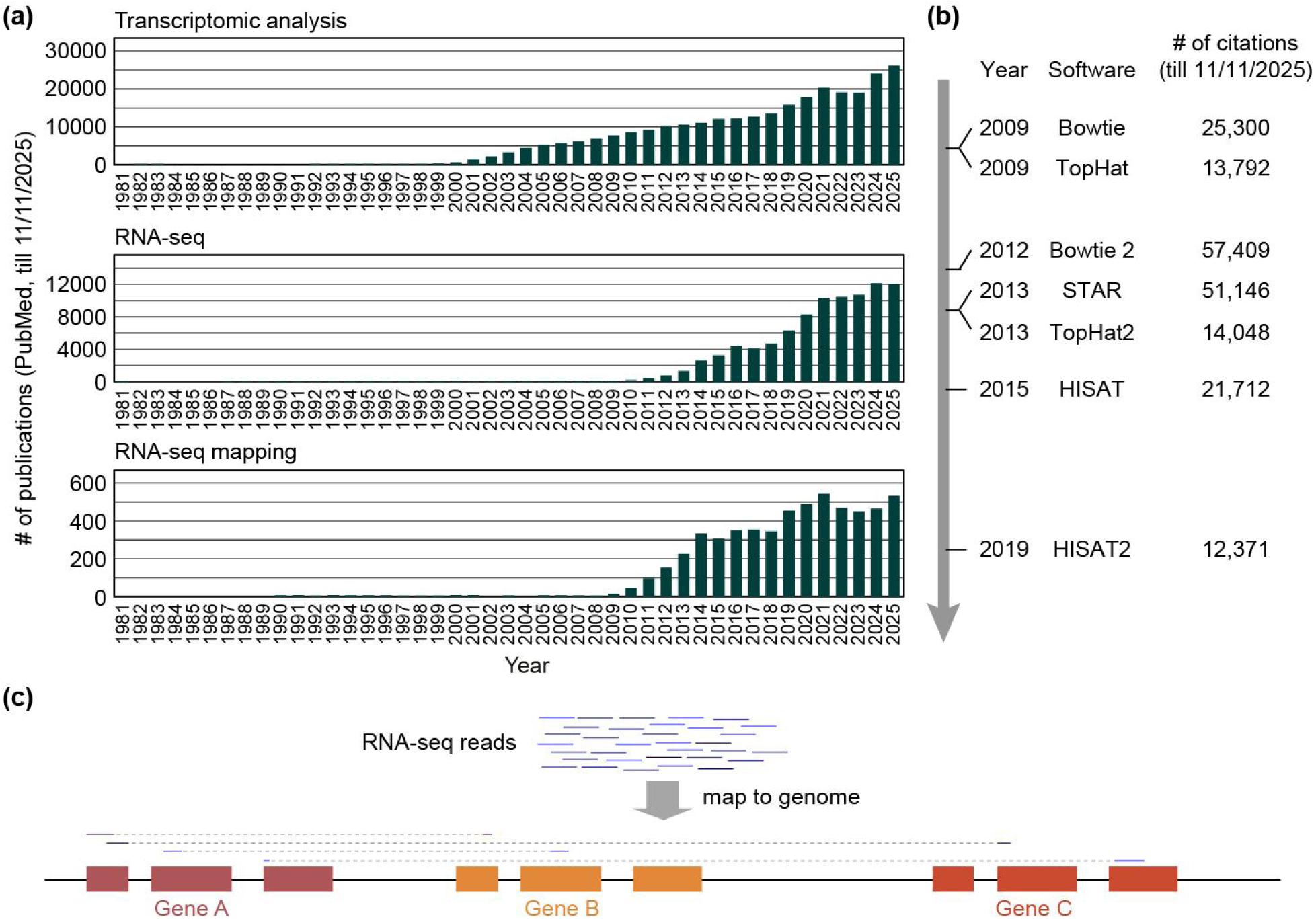
Trends in transcriptomic research since the 1980s and a representative read-mapping pitfall for tandem gene duplicates under the genome-guided mapping strategy. **(a)** Annual number of publications in the PubMed database related to the keywords “transcriptomic analysis”, “RNA-seq” and “RNA-seq mapping” from 1981 to 2025 (searched on Nov 11, 2025). **(b)** Citation counts of the classic alignment tools in the Google Scholar database (searched on Nov 11, 2025). Notably, TopHat was built on Bowtie v1, whereas TopHat2 was built on Bowtie v2. **(c)** Schematic illustration of potential misalignments in G-guided read mapping. Owing to high sequence similarity between tandem duplicates (gene A, B, and C), reads originating from one gene locus can be erroneously mapped across the homologous regions of another copy, resulting in chimeric alignments.

TopHat/TopHat2, as one of the earliest widely adopted splice-aware alignment tools, integrates Bowtie-based read mapping with splice junction prediction, enabling the identification of splicing events in eukaryotic transcriptomes (Kim *et al*., 2013; Trapnell *et al*., 2009). HISAT/HISAT2 is an upgraded alignment algorithm optimized for speed, memory efficiency, and mapping accuracy. By adopting a hierarchical indexing strategy for genome sequences, it significantly reduces memory consumption while maintaining high sensitivity in detecting splice sites and mapping short and long reads (Kim *et al*., 2015, 2019). Compared with TopHat2, HISAT2 exhibits superior computational efficiency and better compatibility with high-throughput sequencing datasets, making it suitable for large-scale transcriptomic cohort analysis. Owing to these advantages, HISAT2 has gradually supplanted TopHat2, which has received little maintenance and support since 2016 (as stated on the TopHat homepage). STAR stands out for its exceptional accuracy and robustness in splice junction mapping (Dobin *et al*., 2013). It employs a two-pass alignment strategy to identify split reads and resolve ambiguous mapping events, achieving the highest precision in detecting complex alternative splicing patterns and novel transcript isoforms. Nevertheless, STAR requires substantial memory resources, which limits its applicability in computing environments with limited resources (Almodaresi *et al*., 2021).

However, due to the pervasive duplicated sequences in eukaryotic genomes, a considerable number of short RNA-seq reads fail to map uniquely to target loci (Kahles *et al*., 2016). This results in ambiguous multi-mapping signals that severely compromise mapping accuracy and downstream differential expression analysis, alternative splicing identification, and transcript quantification.

Specifically, the high sequence similarity characteristic of recent tandem duplications further exacerbates the issues. In our previous project (Jia *et al*., 2026), we found that the default settings of these RNA-seq mapping tools—aligning RNA-seq reads directly to the genome—often failed to accurately quantify the expression of tandem gene duplicates. This is mainly due to the misalignment of individual reads among multiple tandem gene duplicates, as illustrated in **Fig. 1c**. Considering that tandem gene duplicates are prevalent in plant genomes (Panchy *et al*., 2016), and tend to be involved in responses to environmental stimuli and secondary metabolic processes (Birchler and Yang, 2022; Hanada *et al*., 2008; Song *et al*., 2025; Wang *et al*., 2018), given their high sequence similarity, accurate alignment of RNA-seq reads to these genes is particularly critical.

Alternative mapping strategies, such as transcriptome-guided (T-guided) alignment, may help mitigate these quantification inaccuracies. TopHap2 allows an optional mapping strategy: reads are first aligned against transcriptome, then the unaligned reads are aligned against genome (Kim *et al*., 2013). This strategy was initially designed to avoid erroneous alignment of reads to pseudogenes and allow the accurate alignment of reads that cover microexons, non-canonical (such as GC-AG) splice sites, and other “unusual” features of eukaryotic transcriptomes (Kim *et al*., 2013). STAR also offers an option to obtain information about possible splice junction loci from annotation files to avoid aligning reads to pseudogenes (Dobin *et al*., 2013). HISAT2 allows mapping of RNA-seq reads to either the genome index or transcriptome index, but not both. Previous studies have compared the mapping performance among these aligners with others, in terms of mapping rates, computational speed and memory requirements, basewise accuracy, etc (Baruzzo *et al*., 2017; Engström *et al*., 2013; Musich *et al*., 2021; Raplee *et al*., 2019). The alignment performance has also been compared between mapping strategies with or without using annotation as a guide (Baruzzo *et al*., 2017; Engström *et al*., 2013). While these studies have demonstrated that annotation-guided mapping improves alignment quality, most researchers in either the animal or plant field still employ the default genome-guided (G-guided) approach, as shown in **Table S1**. In addition, it remains largely unclear how often RNA-seq reads are misaligned to tandem gene duplicates under default settings (G-guided), and to what degree T-guided strategies can resolve this issue.

In this study, we systematically compared the performance of three classic aligners (TopHat2, HISAT2, and STAR) under two mapping strategies for RNA-seq analysis in six representative plant species—*Arabidopsis thaliana* and five major crops (tomato, potato, grapevine, rice, and maize)—with a particular focus on tandem gene duplicates. Although TopHat2 has not been actively maintained since 2016, we evaluated it here because a substantial number of researchers still use it in transcriptome analysis (**Table S1**). We visually inspected read alignments from six aligner–strategy combinations using real RNA-seq data. Then we generated simulated RNA-seq data based on the expression distribution of real datasets and benchmarked the three aligners in terms of mapping accuracy and computational resource usage, comparing their performance under both G-guided and T-guided mapping strategies. Furthermore, we evaluated the downstream consequences of different mapping strategies, focusing on differential gene expression calls and subsequent Gene Ontology (GO) enrichment results. Our work highlights the previously overlooked issue of RNA-seq read mapping, offering valuable insights for standardizing RNA-seq analysis pipelines.

## Materials and Methods

### Data sources and reference genomes

Genome sequences and the annotation information were downloaded from The Arabidopsis Information Resource (TAIR, https://www.arabidopsis.org/) for *Arabidopsis thaliana* (genome: TAIR 10, annotation: Araport11), Sol Genomics Network (SGN, https://solgenomics.net/) for cultivated tomato (*Solanum lycopersicum*, SL5.0), Ensembl Plants (https://plants.ensembl.org/) for grapevine (*Vitis vinifera*, ASM3070453v1), potato (*Solanum tuberosum*, SolTub3.0) and maize (*Zea mays*, Zm-B73-REFERENCE-NAM-5.0), The Rice Annotation Project Database (RAP-DB, https://rapdb.dna.naro.go.jp/) for rice (*Oryza sativa*, IRGSP-1.0). Raw RNA-seq data were downloaded from The National Center for Biotechnology Information (NCBI) or National Genomics Data Center (NGDC) using accession IDs listed in **Table S2,S3**.

### RNA-seq read mapping

Raw RNA-seq data were processed with Trimmomatic-0.39 (Bolger *et al*., 2014) to remove adapters and discard reads containing ambiguous nucleotides (Ns) or those with low average quality scores, with parameters ILLUMINACLIP:TruSeq3-PE.fa:2:30:10 LEADING:3 TRAILING:3 SLIDINGWINDOW:4:15 MINLEN:36. Prior to read mapping, we built both genome and transcriptome indices for each aligner. For TopHat2 (v2.1.1) (Kim *et al*., 2013), the genome index was generated using bowtie2-build (Langmead and Salzberg, 2012) and the transcriptome index was built with the -G parameter. For HISAT2 (v2.2.1), the genome index was built using hisat2-build. The transcriptome-aware index was constructed by extracting splice sites and exons from the annotation file with the provided Python scripts (extract_exons.py and extract_splice_sites.py, https://github.com/DaehwanKimLab/hisat2/, accessed on Aug 18, 2026), followed by hisat2-build with the --ss and --exon options (Kim *et al*., 2019). For STAR (v2.7.11b), we generated the genome index using --genomeFastaFiles, and with the --sjdbGTFfile option to incorporate splice junctions(Dobin *et al*., 2013). To ensure comparability across the three aligners, we retained only one alignment (randomly selected) for reads that mapped to multiple loci with identical best scores, using the following parameters: --max-multihits 1 (TopHat2), -k 1 (HISAT2), and --outFilterMultimapNmax 1 (STAR).

### Gene expression quantification and differential expression analysis

HTSeq-count (v2.0.5) (Putri *et al*., 2022) was used to count reads mapped to each gene. HTSeq-count provides three modes (--mode) to handle reads that overlap with genes: 1) union (default), a read is assigned to a gene if it overlaps that gene at any position; 2) intersection-strict, a read is assigned to a gene only if every base of the read overlaps that gene; 3) intersection-nonempty, it excludes unannotated positions (e.g., intronic gaps) from the intersection and assigns it to a gene only when all annotated bases unanimously support the same single gene. When a read spans multiple tandemly duplicated genes (**Fig. 1c**), it is assigned to none of these genes under the intersection-strict and intersection-nonempty modes, but is assigned to all matched genes under the union mode. To retain these multi-mapping reads, we employed the default union mode for our analysis.

If a read (or read pair) is assignable to multiple genes, HTSeq-count offers the --nonunique option to handle it. With --nonunique none (the default), the read is discarded and not counted for any gene. Under --nonunique all, the read is counted for all genes. With --nonunique fraction, the read count is divided equally among all assigned genes. Finally, with --nonunique random, the read is assigned to a single gene chosen uniformly at random from the overlapping candidates. To evaluate the impact of the assignment strategies, we compared the quantification results obtained from these four settings.

Transcripts per million (TPM) were calculated by dividing the read counts by the effective transcript length in kilobases, scaling by the sum of all scaled counts, and multiplying by 10^6^. Fold changes (FC) in gene expression levels between samples were calculated using DESeq2 (Love *et ^a^l.*, 2014^)^. Genes with |log_2_(Fold Change)| > 1 and adjusted (Benjamini-Hochberg) *p*-values < 0.05 were considered differentially expressed between samples.

### Simulation of benchmark RNA-seq data

To generate benchmark RNA-seq data that faithfully recapitulate the expression profiles of real experiments, we simulated paired-end 150-bp reads using a custom Python script (see **Data availability**). The script was written by Deepseek under supervision of the authors. The simulation was based on the *Arabidopsis* TAIR10 reference genome and Araport11 annotation (GFF file). For each annotated gene, we selected the longest transcript and extracted its spliced exonic sequence. The read counts obtained from three real RNA-seq datasets served as the ground-truth expression matrices. For each gene, the exact number of paired-end fragments were randomly sampled uniformly along the transcript sequence. Fragment lengths were drawn from a normal distribution with a mean of 300 bp and a standard deviation of 50 bp. Additionally, for each gene, we compared the true read count and the resulting count based on the simulated RNA-seq data, in order to evaluate the mapping accuracy of three aligners under two different mapping strategies.

### Identification of tandem gene duplicates

To identify tandem gene duplicates in the genomes, we employed MCScanX-transposed (Wang *et al*., 2013). Top five best blastp matches within species and across species were obtained first. These data, along with the corresponding GFF files, were subsequently processed by MCScanX-transposed.pl (with default settings) to categorize gene duplicates into tandem, proximal, segmental, and dispersed types.

### GO enrichment analysis

GO annotations for genes in all six plant species were obtained from the same databases that provided the corresponding genome assemblies and annotations. GO enrichment analysis was conducted using R package clusterProfiler (v4.0) (Xu *et al*., 2024). GO terms with adjusted *p*-values < 0.05 were considered significantly enriched for a set of differentially expressed genes (DEGs).

## Results

### Substantial misalignment of RNA-seq reads at tandem gene duplicates under genome-guided mapping

To compare the performance of G-guided and T-guided mapping strategies across three commonly used aligners, we first employed the root phloem and companion cell RNA-seq data (SRA accession: SRR3664372) from Li *et al*., (2016), where transcriptomes of different root cell types in *A. thaliana* were sequenced. To quantify the number of genes showing differential expression levels (TPMs) between the two mapping strategies, we generated scatter plots of TPM_G_ (TPM values called based on G-guided mapping strategy) vs TPM_T_. For each gene, we calculated the absolute distance from its data point to the diagonal line (where TPM_G_ = TPM_T_). We then computed *z*-scores for all absolute distances across the entire gene set. Genes with an z-score > 3 (corresponding to a two-sided *p*-value of 2.7 × 10^−3^) were considered to have significantly different TPMs between the two strategies.

When the --nonunique all parameter was used in HTSeq-count for expression quantification (**Methods**), 0.85% (TopHat2), 0.35% (HISAT2), and 0.54% (STAR) of all 27,106 *Arabidopsis* genes showed significantly different TPM values between the two mapping strategies (**Fig. 2a**). Among these genes, only six genes were detected by all three aligners (**Fig. 2b**). In addition, the number of genes with TPM_T_ < TPM_G_ exceeded that with TPM_T_ > TPM_G_. To elucidate the possible sources of quantification bias, we visually examined RNA-seq read alignments across related genes in Integrative Genomics Viewer (IGV) (Robinson *et al*., 2023). For *AT1G73280* (*serine carboxypeptidase-like 3*), which had TPM_T_ > TPM_G_ in TopHat2, the T-guided strategy produced clean alignments, whereas the G-guided strategy generated misaligned reads spanning between *AT1G73280* and its tandem duplicate *AT1G73290* (*serine carboxypeptidase-like 6*) (**Fig. 2c**). Conversely, for *AT1G60690* (encoding a NAD(P)-linked oxidoreductase superfamily protein), which had TPM_T_ < TPM_G_ in TopHat2, the T-guided strategy yielded reasonable alignments, but the G-guided strategy misaligned partial reads from its tandem duplicates *AT1G60680* and *AT1G60710* to *AT1G60690*, thereby inflating its expression estimate (**Fig. 2c**).

**Fig. 2.**
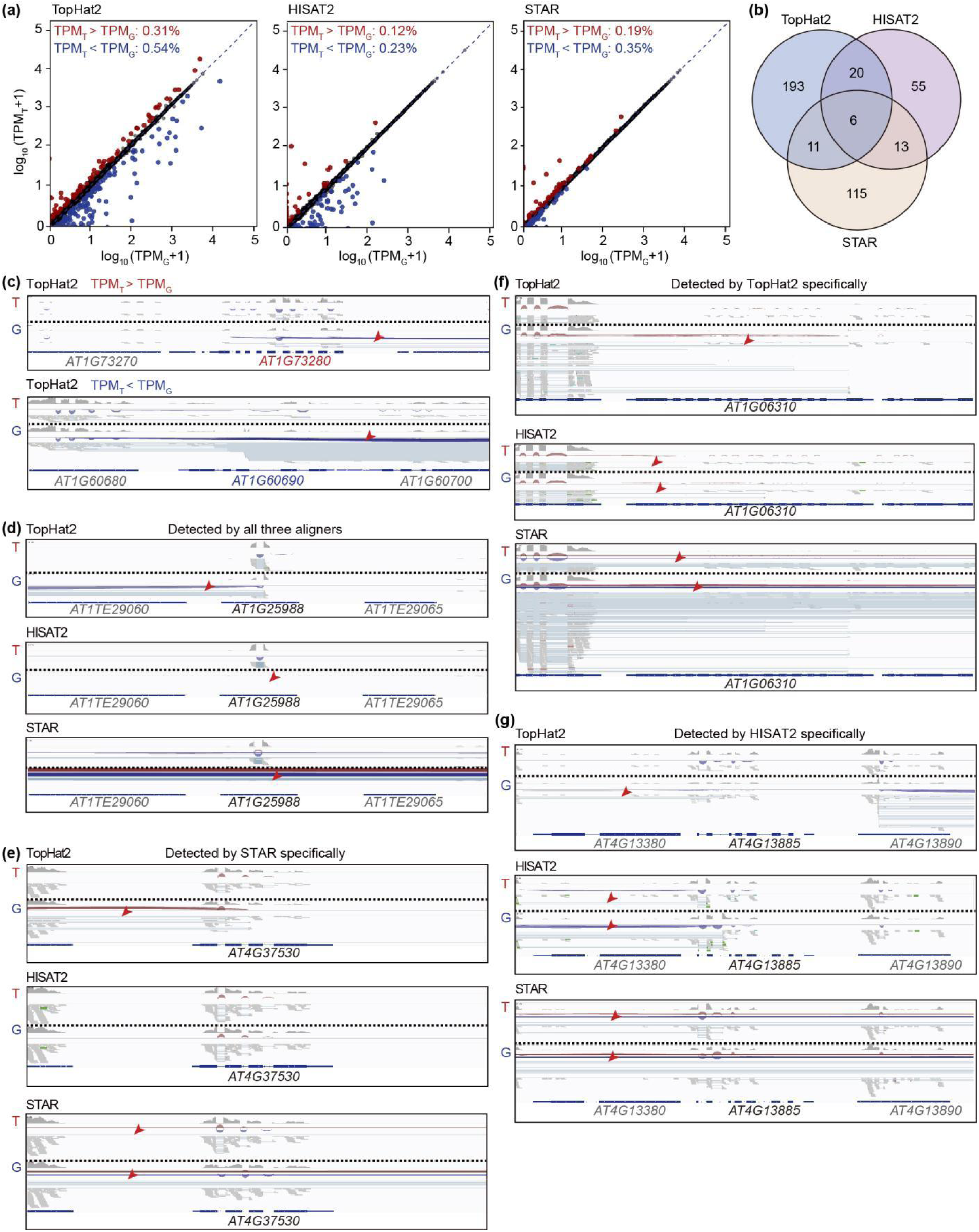
Comparison of gene expression levels derived from genome-guided vs transcritome-guided read-mapping strategies. (**a**) Scatter plots comparing TPM values between G-guided and T-guided strategies for three aligners (TopHat2, HISAT2, and STAR). Red dots indicate genes with significantly higher TPM values in T-guided than in G-guided mapping (TPM_T_ > TPM_G_, *z*-score of the absolute distance to the diagonal > 3; see **Methods**), while blue dots indicate genes with TPM_T_ < TPM_G_. (**b**) Venn diagram showing the overlap of genes that exhibited significant TPM differences between the two strategies across the three aligners. (**c**) Read alignment visualizations for genes *AT1G73280* and *AT1G60690*, which showed TPM_T_ > TPM_G_ and TPM_T_ < TPM_G_, respectively, in TopHat2 analyses. (**d–g**) Read alignment visualization for genes which displayed significant TPM differences between the two strategies across all three aligners (**d**), STAR only (**e**), TopHat2 only (**f**), and HISAT2 only (**g**). Red arrows highlight misaligned reads or alignment errors.

For *AT1G25988* (encoding a coiled-coil protein) that was detected by all three aligners, all three T-guided mapping efforts produced reasonable alignments (**Fig. 2d**). In contrast, the G-guided strategy yielded alignment errors that varied across aligners: reads spanning *AT1G25988* and its tandem duplicate *AT1G25682* were misaligned in TopHat2, reads were completely absent in HISAT2 and severely reduced in STAR. For *AT4G37530* (encoding a Peroxidase superfamily protein)—a gene specifically detected by STAR, G-guided TopHat2 and STAR yielded alignment errors (**Fig. 2e**), whereas T-guided TopHat2 corrected all the errors, and T-guided STAR corrected most of them. For *AT1G06310* (encoding a putative acyl-CoA oxidase), which was specifically detected by TopHat2, the G-guided strategy inflated its expression estimate by misaligning partial reads from its tandem duplicate *AT1G06290* (**Fig. 2f**). By contrast, T-guided TopHat2 corrected all such errors, whereas T-guided HISAT2 and STAR corrected most, but not all, of them. Similarly, for *AT4G13885* (encoding a Polynucleotidyl transferase), T-guided TopHat2 corrected G-guided errors, whereas T-guided HISAT and STAR corrected most, but not all (**Fig. 2g**).

Taken together, these results reveal systematic alignment biases between the two mapping strategies and across the three aligners. T-guided TopHat2 outperformed the other configurations in alignment accuracy for the RNA-seq data examined here. T-guided HISAT2 and STAR mitigated the errors seen in their G-guided versions, though some misalignments persisted.

Importantly, the lower number of genes with substantial TPM differences between G-guided and T-guided strategies and the reduced dispersion observed with HISAT2 and STAR, compared with TopHat2 (**Fig. 2a,b**), do not necessarily reflect superior alignment quality, at least for these tandem gene duplicates discussed above.

### Impact of multi-mapping read assignment on expression quantification

In the above section, we adopted --nonunique=all in HTSeq-count to handle multi-mapping reads during expression quantification. We found that the number of genes showing TPM differences between the two mapping strategies varied with different --nonunique parameter settings in HTSeq-count (**Fig. 3a**, only TopHat2 results were presented). Compared with the -- nonunique=all setting (used above), the default --nonunique=none option—which is also commonly adopted in the community—yielded significantly lower expression (z-score > 3) for 1.81% of genes (**Fig. 3b**, for clarity, only results from T-guided TopHat2 were presented). By contrast, the TPM differences relative to --nonunique=all were markedly reduced when using either the --nonunique=fraction (2.96%) or --nonunique=random (2.67%) option, although the proportion of genes increased (**Fig. 3b**).

**Fig. 3.**
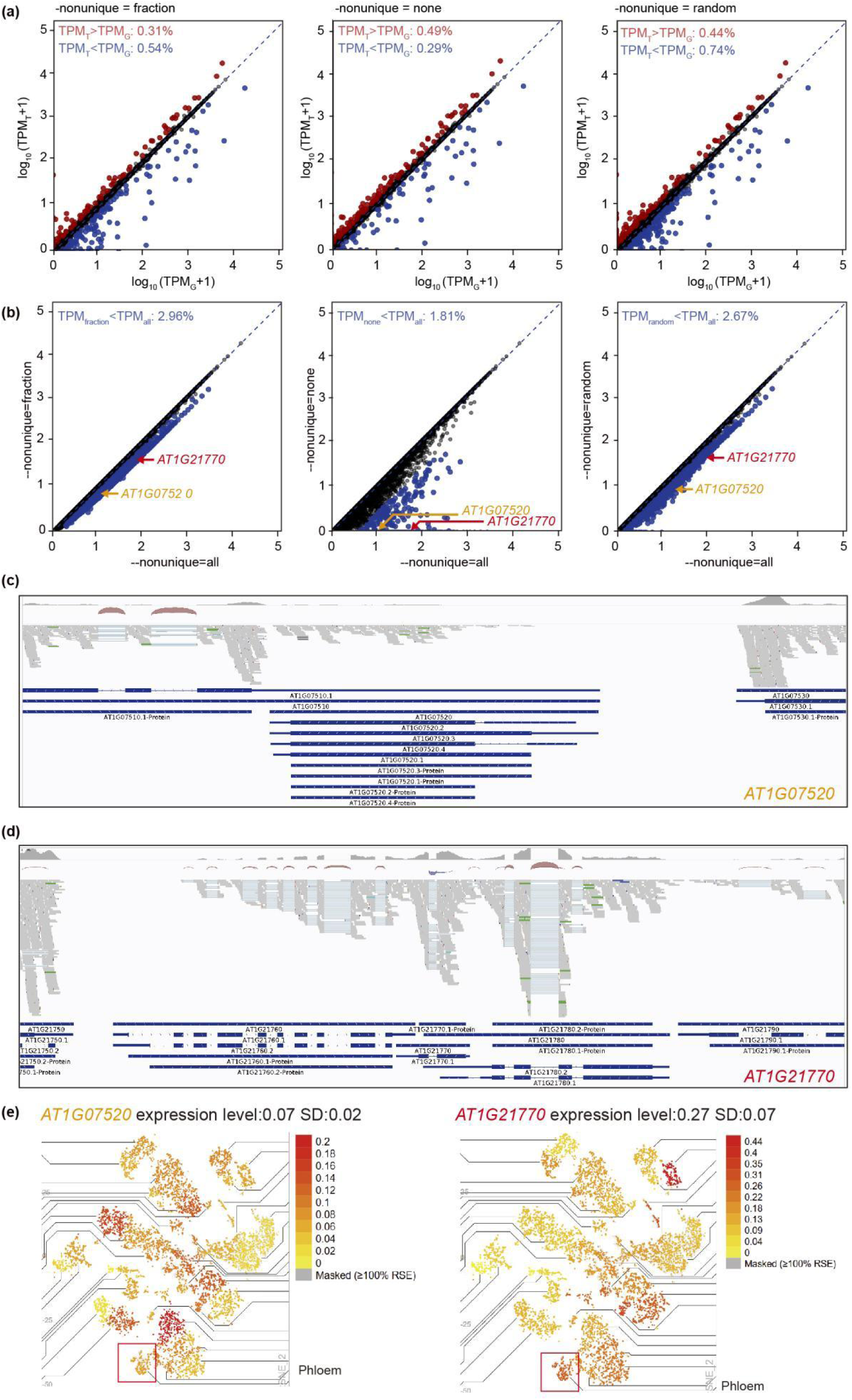
Influence of the HTSeq-count --nonunique parameter on expression quantification. (**a**) Scatter plots comparing TPM values obtained from G-guided vs T-guided TopHat2 alignments, using selected -- nonunique settings (fraction, none, and random). Red and blue dots indicate genes with TPM_T_ > TPM_G_ and TPM_T_ < TPM_G_, respectively (*z*-score of the absolute distance to the diagonal > 3). (**b**) Scatter plots comparing TPM values quantified with different --nonunique parameters specifically in T-guided TopHat2 analyses. Blue dots denote genes for which the --nonunique all setting produced significantly higher TPM values than the other options tested; data points for the highlighted genes *AT1G07520* and *AT1G21770* are labeled. (**c,d**) Read alignment visualization for *AT1G07520* (**c**) and *AT1G21770* (**d**), which were assigned zero TPM values with --nonunique none (default) but nonzero TPM values with other three settings, illustrating the direct impact of multi-mapping read handling on quantification. (**e**) Expression patterns of *AT1G07520* and *AT1G21770* in phloem tissues, retrieved from the TAIR BAR eFP Browser.

To investigate which parameter setting yielded more accurate expression quantification, we examined read alignments for two genes—*AT1G07520* (encoding a GRAS family transcription factor) and *AT1G21770* (encoding an Acyl-CoA N-acyltransferases (NAT) superfamily protein)—that had zero expression under the --nonunique=none setting but showed non-zero expression under the other three parameter settings. Both genes showed clean read alignments without evident errors, indicating active expression (**Fig. 3c,d**), which was also supported by single-cell RNA-seq data from a previous study (Ryu *et al*., 2019), as displayed in the TAIR BAR eFP Browser (**Fig. 3e**). Both genes overlap with neighboring gene annotations, which led to their reads being classified as multi-mapping and discarded under --nonunique=none. Given that 24.42% (6,619 out of 27,106) of genes in the *Arabidopsis* genome overlap with other gene annotations, the default HTSeq-count setting inevitably yields lower expression estimates than actual levels for a substantial proportion of genes. However, none of the --nonunique=all, -- nonunique=fraction, or --nonunique=random options can properly resolve such cases. Instead, accurate quantification would benefit from more precise gene boundary definition and enhanced annotation quality, achievable via targeted experiments or full-length transcriptome data.

Nevertheless, for subsequent analysis, we adopted the --nonunique=fraction option for expression quantification, as --nonunique=all would overestimate total read counts, whereas -- nonunique=random could compromise analysis reproducibility.

### Performance of three aligners under two mapping strategies on simulated RNA-seq data

To directly compare the mapping accuracy of three aligners under two mapping strategies, we simulated RNA-seq reads based on the real read counts obtained from three RNA-seq datasets (**Fig. 4**) in a T-guided TopHat2 analysis with the --nonunique fraction option, using the *Arabidopsis* genome sequence and annotation files (**Methods**). The simulated reads were then mapped to the transcriptome and/or genome, and read counts were calculated for each gene. With respect to computational resource usage, consistent with previous studies (Dobin *et al*., 2013; Kim *et al*., 2015), TopHat2 was the slowest aligner, whereas STAR was the fastest but required substantially more memory than the others for large datasets (**Fig. 4a,b**). Regarding mapping accuracy, the T-guided strategies generally outperformed the G-guided ones, yielding higher concordant mapping rates and improved accuracy, as reflected by smaller mean squared errors (MSE) between the true reference counts and the estimated counts (**Fig. 4c,d**).

**Fig. 4.**
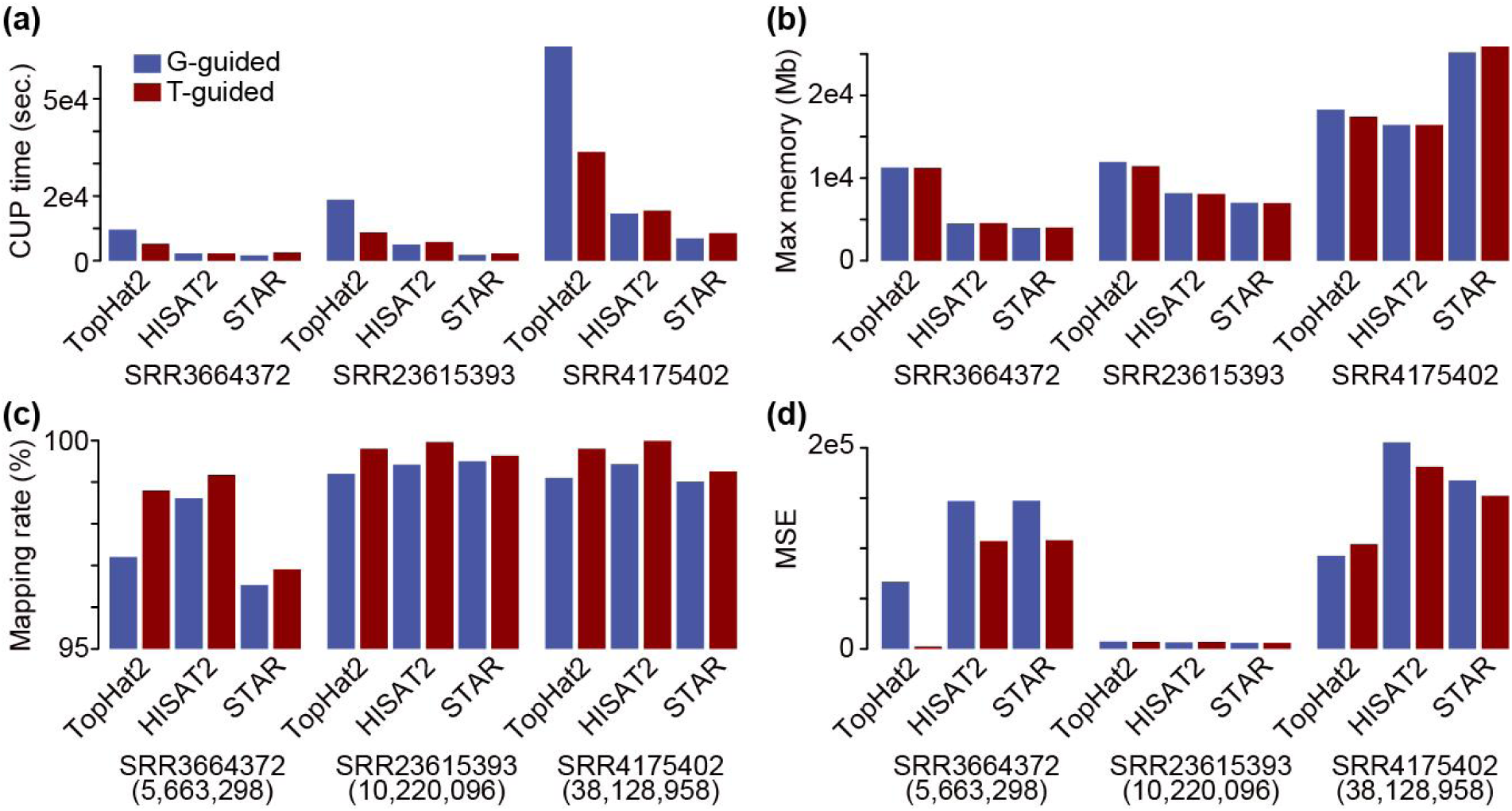
Mapping statistics of simulated RNA-seq data across three aligners under two strategies. (**a**) CPU time used. (**b**) Max memory used. (**c**) Concordant mapping rate. (**d**) Mean squared error (MSE) between real and simulated read counts. The total number of simulated reads is indicated in parentheses.

With regard to the aligners, the optimal choice varied across datasets. For the simulated data derived from SRR3664372 (root phloem and companion cell), T-guided TopHat2 achieved the highest accuracy. For SRR23615393 (leaves treated with heat-drought stress), T-guided STAR gave the smallest MSE, although the MSE values from all six aligner–strategy combinations were relatively similar. For SRR4175402 (leaves treated with 10 mM MgCl_2_), G-guided TopHat2 performed best, with only a marginal advantage over its T-guided counterpart. Notably, mapping rate did not always correlate with mapping accuracy; the Spearman’s correlation coefficient (ρ) between mapping rate and MSE was -0.16 (*p*-value = 0.52). Given these observations and findings discussed above, we focused on TopHat2 in the subsequent analyses to simplify the presentation.

### Overrepresentation of alignment bias between mapping strategies in tandem gene duplicates

Given that the genes displaying alignment errors upon IGV inspection in the above section were predominantly located in tandemly duplicated arrays, we asked how frequently such errors originated from tandem gene duplicates. We classified genes into four different categories: tandem (consecutive to each other)/proximal (separated by a few non-paralogous genes) duplicates, segmental duplicates (collinear gene pairs), transposed (duplicated from an ancestral locus to a novel locus) duplicates, and the remaining dispersed duplicates (**Methods**). For simplicity, we collectively referred to tandem and proximal duplicates as tandem duplicates hereafter. In *Arabidopsis* (**Fig. 5a**), potato (**Fig. 5c**) and maize (**Fig. 5f**), genes with TPM_T_ < TPM_G_ were significantly enriched for tandem duplicates in all samples examined (odds ratio > 4, *p*-values < 0.001), while those with TPM_T_ > TPM_G_ were significantly enriched for tandem duplicates only in 0–4 samples. In tomato (**Fig. 5b**), grapevine (**Fig. 5d**) and rice (**Fig. 5e**), both TPM_T_ < TPM_G_ and TPM_T_ > TPM_G_ categories were significantly enriched for tandem duplicates in almost all samples. These results further confirmed the impact of mapping strategy choice on expression quantification of tandem gene duplicates. Across the six plant species examined here, tandemly duplicated genes generally showed a functional bias toward defense and stress responses, detoxification and secondary metabolism (**Fig. 5a–f**). This pattern aligns well with previous findings in diverse angiosperms (Hanada *et al*., 2008; Song *et al*., 2025; Yang *et al*., 2025), reinforcing the notion that tandem duplication is a key driver of adaptive and metabolic innovation. Therefore, precise expression quantification of tandemly duplicated genes is essential for plant researches aimed at elucidating these biological processes.

**Fig. 5.**
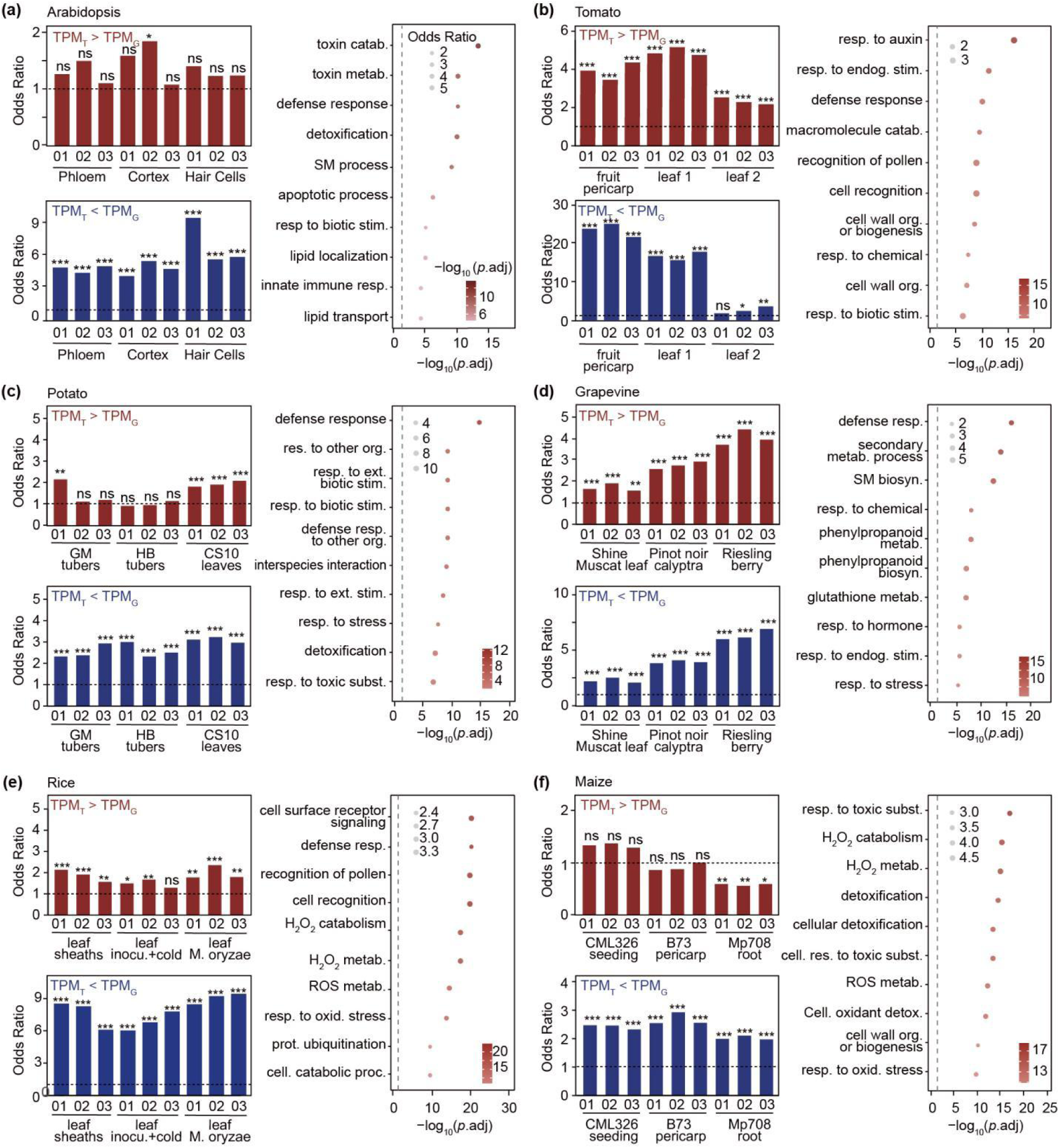
Enrichment of genes with different TPM values between two strategies for tandem gene duplicates. (**a–f**) Enrichment (Fisher’s exact test) of genes with differential TPM values between the two strategies within tandem/proximal duplicates (left panel) and enriched GO terms of tandem/proximal genes (right panel) in *Arabidopsis* (**a**), tomato (**b**), potato (**c**), grapevine (**d**), rice (**e**) and maize (**f**). ns: not significant. *, **, and *** indicate *p*-values < 0.05, 0.01, and 0.001, respectively. Sample information and the corresponding SRA data, see **Supplementary Table S2**.

### Impact of mapping strategies on functional enrichment

To assess the influence of mapping strategy choice on DEG identification and downstream functional enrichment, we downloaded RNA-seq data from paired samples generated from experiments designed to investigate the biological processes represented by these tandem-duplicate-enriched GO terms (**Table S3**). The proportion of DEGs and enriched GO terms that differed between the two mapping strategies varied across datasets, ranging from 1.95% to 20.83% for DEGs and 0% to 100% for GO terms (**Fig. 6a**, **Table S4**). Most GO terms that differed in significance between the two mapping strategies had adjusted *p*-values close to the threshold (0.05); thus, a small fluctuation in the number of identified DEGs could shift the significance of the associated GO terms (e.g., those in **Fig. 6b–e**). In contrast, the GO terms in **Fig. 6f,g** exhibited remarkably different *p*-values between the two strategies, suggesting fundamental differences in how the two strategies quantify gene expression, which consequently leads to divergent functional interpretations.

**Fig. 6.**
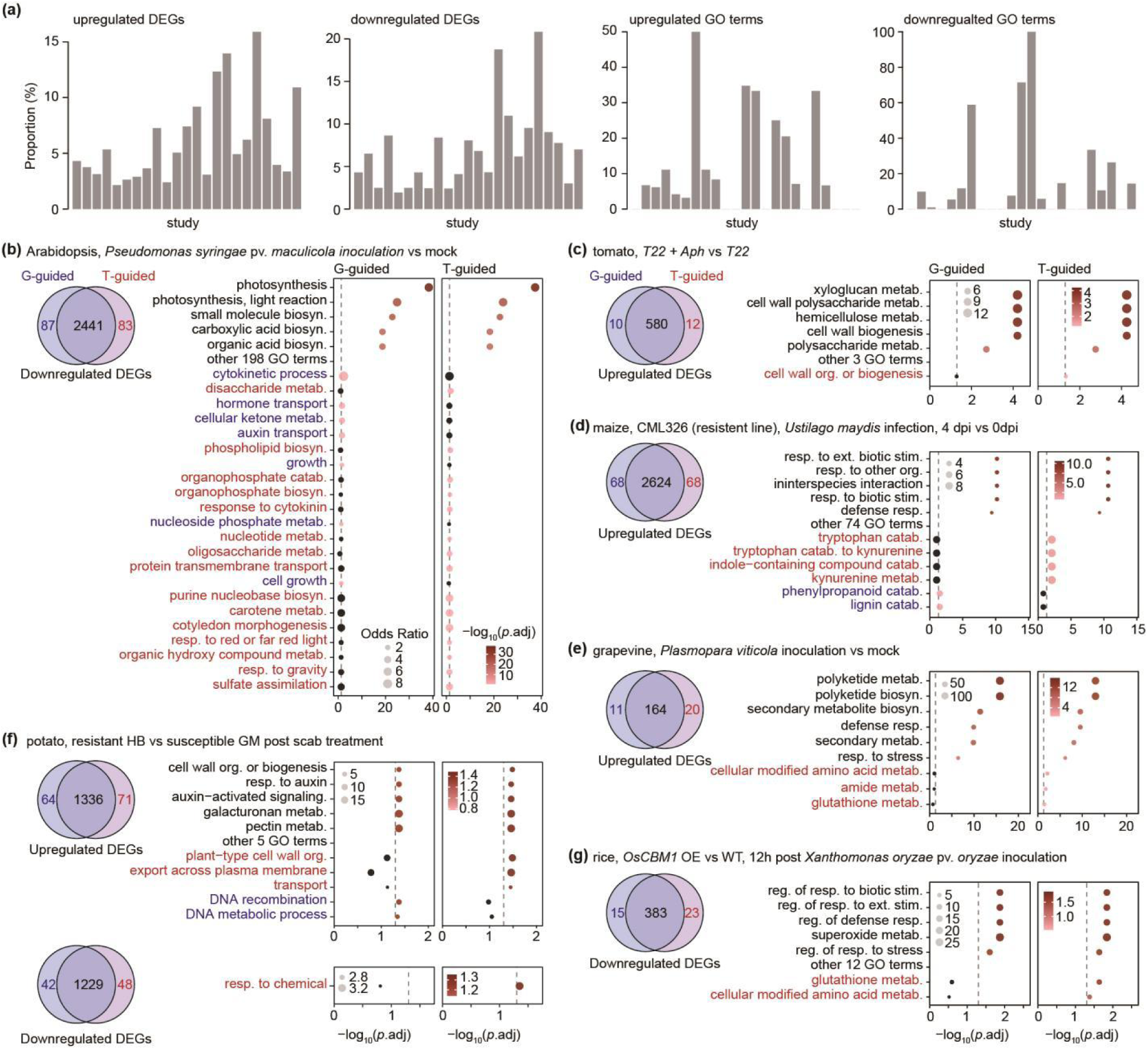
Impact of mapping strategies on DEG identification and GO enrichment analysis. (**a**) Proportions of unique DEGs and enriched GO terms identified by the two mapping strategies. (**b**) Downregulated DEGs in *Arabidopsis* seedlings at 48 h after inoculation with *Pseudomonas syringae* pv. *maculicola* vs mock-treated. (**c**) Upregulated DEGs in tomato leaves treated with *Trichoderma harzianum* (strain T22) plus the *aphid Macrosiphum euphorbiae* vs *T. harzianum* alone. (**d**) Upregulated DEGs in resistant maize line CML326 at 4 vs 0 days post inoculation (dpi) with *Ustilago maydis*. (**e**) Upregulated DEGs in grapevine leaves at 1 dpi with *Plasmopara viticola* vs mock-treated. (**f**) Up- and downregulated DEGs between scab-resistant potato cultivar Hindenburg (HB) and susceptible Green Mountain (GM) after *Streptomyces scabies* incubation. (**g**) Downregulated DEGs in rice *OsCBM1* overexpression line vs WT at 12 h post *Xanthomonas oryzae* pv. *oryzae* inoculation. GO terms enriched only under T-guided mapping are in red; those only under G-guided mapping are in blue. See **Supplementary Table S3** for sample information and SRA data.

For instance, in the comparison between scab-resistant potato cultivar Hindenburg (HB) vs the susceptible cultivar Green Mountain (GM) after scab (*Streptomyces scabies*) infection (Fofana et al., 2020) (**Fig. 6f**), GO terms “plant−type cell wall organization”, “export across plasma membrane”, and “transport” were enriched for upregulated DEGs specifically by the T-guided strategy, whereas “DNA recombination” and “DNA metabolic process” were exclusively detected by the G-guided approach. In addition, the GO term “response to chemical” was specifically enriched for downregulated DEGs in HB compared with GM, and this enrichment was only captured by the T-guided method. These results collectively demonstrate that the T-guided mapping strategy offers superior sensitivity in resolving genotype-specific transcriptional reprogramming during the defense response. Specifically, it effectively captured the activation of cell wall remodeling and transport processes in the resistant cultivar—both of which are hallmarks of active defense—while also revealing the suppression of chemical response pathways that may represent a susceptibility-associated program attenuated in the resistant genotype (Bhandari et al., 2025; Khare et al., 2017; Mohr and Cahill, 2007).

*OsCBM1*, encoding a carbohydrate-binding malectin/malectin-like domain-containing protein in rice, has been reported to contribute to drought-stress tolerance by enhancing reactive oxygen species (ROS) burst (Jing *et al*., 2021). The *OsCBM1* OE line was shown to exhibit heightened resistance to bacterial blight compared with WT (Bie *et al*., 2025). The GO term “glutathione metabolic process” was significantly enriched among downregulated DEGs in the *OsCBM1* OE line specifically in the T-guided analysis (**Fig. 6g**). This finding aligns with the enhanced ROS burst phenotype, as the repression of glutathione metabolism would reduce the cellular capacity for ROS detoxification (Kozieł *et al*., 2024; Noctor and Foyer, 1998), thereby reinforcing oxidative stress. In contrast, glutathione metabolism was significantly upregulated in a resistant grapevine cultivar (Shine Muscat) inoculated with *Plasmopara viticola* (**Fig. 6e**, Shimizu and Suzaki, 2020), which likely serves to maintain redox homeostasis and fine-tune the ROS burst, consistent with previous findings (Liu *et al*., 2021). Similar to glutathione metabolism, cellular modified amino acid metabolism was also downregulated in *OsCBM1* OE lines but upregulated during pathogen attack (**Fig. 6e,g**). This contrast suggests that while pathogen infection actively mobilizes amino acid-derived antioxidant and detoxification machinery, *OsCBM1* overexpression compromises this biochemical defense capacity, thereby exacerbating oxidative stress. The detection of these processes exclusively by the T-guided strategy further underscores its sensitivity in uncovering functionally relevant transcriptional events that may contribute to the observed stress-tolerant phenotypes.

For GO terms showing borderline significance under both mapping strategies, G-guided alignment may also lead to misleading biological interpretations. For example, upon infection with the biotrophic fungal pathogen *Ustilago maydis*, the maize resistant line CML326 displayed a specific transcriptional reprogramming at 4 days post-inoculation (dpi) compared with 0 dpi (Ruan *et al*., 2021). T-guided and G-guided mapping strategies shared 79 commonly enriched GO terms among the upregulated DEGs, while four additional GO terms were uniquely identified by the T-guided method but with significance level close to the threshold (**Fig. 6d**).

These four terms were “tryptophan catabolic process”, “tryptophan catabolic process to kynurenine”, “indole-containing compound catabolic process”, and “kynurenine metabolic process”, all of which were interconnected and centered on tryptophan degradation and indole compound catabolism. The upregulation of genes involved in these processes likely reflected a defensive strategy to counter *U. maydis* infection. Given that the pathogen itself utilized tryptophan to produce the indole-3-acetic acid (IAA) (Reineke *et al*., 2008), the host’s enhanced catabolism of these substrates may serve to deplete resources critical for fungal pathogenesis and mitigate the accumulation of signaling molecules that the pathogen could exploit.

Taken together, these results collectively demonstrate that the choice of mapping strategy is a critical determinant of downstream functional interpretation. The T-guided approach, by virtue of its superior alignment precision, enabled more reliable DEG identification and, consequently, more biologically meaningful GO enrichment outcomes, underscoring its general applicability and recommending its adoption as a preferred practice in transcriptomic analyses.

## Discussion

Our systematic comparison of G-guided and T-guided mapping strategies across six plant species revealed that the choice of mapping strategy had a profound and previously underappreciated impact on gene expression quantification and downstream functional interpretation. While the importance of alignment accuracy in RNA-seq analysis has long been recognized (Baruzzo *et al*., 2017; Musich *et al*., 2021; Raplee *et al*., 2019; Stark *et al*., 2019), our results demonstrate that this issue is particularly acute for tandemly duplicated genes—a gene category that is disproportionately enriched for functions in defense and stress responses, detoxification, and secondary metabolism.

Baruzzo *et al*. (2017) previously demonstrated that the use of annotation-guided (i.e., T-guided) strategies produced more reliable results than default G-guided approaches (Baruzzo *et al*., 2017). Despite these clear warnings, however, a substantial proportion of RNA-seq studies continue to map reads directly to the genome without leveraging transcriptome annotation (**Table S1**). Our results provide a mechanistic explanation for why this practice can be problematic: in G-guided mapping, reads originating from tandemly duplicated genes are frequently misaligned across paralogs due to their high sequence similarity, leading to systematic quantification biases that propagate through to differential expression analysis and functional enrichment.

Critically, our findings challenge the adequacy of commonly reported alignment metrics such as mapping rates. In our benchmark analyses, mapping rate did not always correlate with mapping accuracy (**Fig. 4**). In addition, HISAT2 and STAR exhibited lower numbers of genes with significantly different expression levels between the two mapping strategies and reduced dispersion compared with TopHat2 (**Fig. 2a**). However, upon visual inspection of read alignments, we found that this apparent “stability” did not necessarily reflect superior alignment accuracy. These observations suggest that mapping rate and related global metrics are insufficient proxies for alignment quality, particularly in complex genomic regions such as tandem arrays. Researchers must therefore evaluate alignment tools, strategies, and parameters in the context of their specific biological question and genomic landscape, rather than relying on global performance statistics.

The implications of our findings extend beyond bulk RNA-seq to the rapidly growing field of single-cell transcriptomics. Current single-cell RNA-seq mapping pipelines are predominantly built upon STAR, which is embedded as the core aligner in popular tools such as 10x Genomics Cell Ranger and STARSolo (Kaminow *et al*., 2021). Our results demonstrate that even STAR, despite its overall robust performance, exhibits alignment errors for tandem gene duplicates under G-guided mapping—errors that were partially but not completely corrected by the T-guided strategy. Given that single-cell data are typically sparser and noisier than bulk data, the impact of such misalignments on cell-type-specific expression estimates may be even more pronounced. There is an urgent need for systematic evaluation of single-cell mapping pipelines regarding their accuracy in quantifying tandem duplicates and other paralogous genes, particularly in plant systems where tandem duplication is a one of the major drivers of adaptive evolution.

Another critical issue highlighted by our study is the dependence of quantification accuracy on gene annotation quality. For genes overlapping with their neighboring genes, reads are inevitably classified as multi-mapping and subsequently discarded in the default HTSeq-count settings—a practice also commonly adopted in the plant research community. None of the --nonunique settings in HTSeq-count could solve this fundamental limitation. Future improvements should therefore prioritize the integration of experimental data, such as full-length transcriptome sequencing (Iso-Seq) or directed experimental validation, to refine gene structure annotations.

With more precise gene boundaries, the fraction of reads classified as multi-mapping would decrease, and the accuracy of quantification—regardless of the mapping strategy employed— would improve substantially.

## Conclusion

Our study systematically demonstrates that the choice of RNA-seq read-mapping strategy— G-guided vs T-guided—had a profound and often underestimated impact on gene expression quantification and downstream functional inference, particularly for tandemly duplicated genes. Across six plant species, we showed that T-guided alignment consistently improved accuracy, especially for TopHat2, although it is no longer actively maintained. Commonly used metrics such as mapping rate were not always suitable proxies for true alignment quality. Importantly, the biases were not merely technical but directly affected the detection of biologically relevant processes—defense, stress responses, secondary metabolism—that were enriched in tandem arrays. Our results further revealed that multi-mapping read handling and annotation precision were critical confounding factors, with default settings frequently discarding genuine expression signals from overlapping genes. We therefore advocate the adoption of T-guided mapping as a recommended practice in plant transcriptomics, alongside community efforts to refine gene annotations. These guidelines will enable more reliable comparative studies and accelerate the functional dissection of adaptive traits governed by tandemly duplicated genes.

## Supplementary Tables

**Table S1** RNA-seq mapping settings in published papers

**Table S2** SRA accession IDs for RNA-seq data used in **Figure 5**

**Table S3** SRA accession IDs for RNA-seq data used in **Figure 6** and **Supplementary Table S4**

**Table S4** Impact of mapping strategies on DEG identification and functional analysis

## Acknowledgments

This work was supported by the National Natural Science Foundation of China (32370241 to PW). During the preparation of this work, the authors used DeepSeek to write the RNA-seq data simulation script and assist with writing refinements. After using this tool/service, the authors reviewed and edited the content as needed and take full responsibility for the content of the published article.

## Author contributions

P.W. and Y.B. conceived and designed the study. Y.J. performed the analysis with assistance from F.W., while P.W. conducted the simulation analysis. Y.J. and P.W. generated figures and tables. P.W. drafted and revised the manuscript, incorporating feedback from all co-authors. All authors read and approved the final manuscript.

## Data availability

All the scripts used in this study are available on GitHub at: https://github.com/peipeiwang6/Manuscript/tree/main/2026_RNA-seq_mapping_strategies.

## Reference

1. Almodaresi, F., Zakeri, M., and Patro, R. (2021) PuffAligner: a fast, efficient and accurate aligner based on the Pufferfish index. Bioinformatics, 37, 4048–4055.

2. Baruzzo, G., Hayer, K.E., Kim, E.J., Di Camillo, B., FitzGerald, G.A., and Grant, G.R. (2017) Simulation-based comprehensive benchmarking of RNA-seq aligners. Nature Methods, 14, 135–139.

3. Bhandari, D.D., Kim, S.-J., and Brandizzi, F. (2025) Fortifying the frontier: cell wall modifications during plant immunity. Current Opinion in Plant Biology, 88, 102816.

4. Bie, S., Xiao, Y., Zhang, L., Liu, Y., He, X., Peng, J., et al. (2025) *Transcriptome analysis reveals the role of* OsCBM1 *in rice defense against* Xanthomonas oryzae pv.oryzae. Biomolecules, 15, 287.

5. Birchler, J.A. and Yang, H. (2022) The multiple fates of gene duplications: Deletion, hypofunctionalization, subfunctionalization, neofunctionalization, dosage balance constraints, and neutral variation. The Plant Cell, 34, 2466–2474.

6. Bolger, A.M., Lohse, M., and Usadel, B. (2014) Trimmomatic: a flexible trimmer for Illumina sequence data. Bioinformatics, 30, 2114–2120.

7. Dobin, A., Davis, C.A., Schlesinger, F., Drenkow, J., Zaleski, C., Jha, S., et al. (2013) *STAR: ultrafast universal RNA-seq aligner*. *Bioinformatics (Oxford*, England), 29, 15–21.

8. Engström, P.G., Steijger, T., Sipos, B., Grant, G.R., Kahles, A., Rätsch, G., et al. (2013) Systematic evaluation of spliced alignment programs for RNA-seq data. Nature Methods, 10, 1185–1191.

9. Fofana, B., Somalraju, A., Fillmore, S., Zaidi, M., Main, D., and Ghose, K. (2020) *Comparative transcriptome expression analysis in susceptible and resistant potato (*Solanum tuberosum*) cultivars to common scab (*Streptomyces scabies*) revealed immune priming responses in the incompatible interaction*. PloS One, 15, e0235018.

10. Hanada, K., Zou, C., Lehti-Shiu, M.D., Shinozaki, K., and Shiu, S.-H. (2008) Importance of lineage-specific expansion of plant tandem duplicates in the adaptive response to environmental stimuli. Plant Physiology, 148, 993–1003.

11. Jia, R., Zuo, X., Zou, L., Wang, L., Lin, L., Wang, Z., et al. (2026) *Haplotype-resolved genome assembly advances genetic understanding of trait diversity in* Phalaenopsis *orchids*. bioRxiv, DOI: 10.64898/2026.02.11.705238.

12. Jing, X.-Q., Li, W.-Q., Zhou, M.-R., Shi, P.-T., Zhang, R., Shalmani, A., et al. (2021) Rice carbohydrate-binding malectin-like protein, OsCBM1, contributes to drought-stress tolerance by participating in NADPH oxidase-Mediated ROS production. Rice, 14, 100.

13. Kahles, A., Behr, J., and Rätsch, G. (2016) MMR: a tool for read multi-mapper resolution. Bioinformatics, 32, 770–772.

14. Kaminow, B., Yunusov, D., and Dobin, A. (2021) STARsolo: accurate, fast and versatile mapping/quantification of single-cell and single-nucleus RNA-seq data. bioRxiv, DOI: 10.1101/2021.05.05.442755.

15. Khare, D., Choi, H., Huh, S.U., Bassin, B., Kim, J., Martinoia, E., et al. (2017) Arabidopsis *ABCG34 contributes to defense against necrotrophic pathogens by mediating the secretion of camalexin*. Proceedings of the National Academy of Sciences of the United States of America, 114, E5712–E5720.

16. Kim, D., Langmead, B., and Salzberg, S.L. (2015) HISAT: a fast spliced aligner with low memory requirements. Nature Methods, 12, 357–360.

17. Kim, D., Paggi, J.M., Park, C., Bennett, C., and Salzberg, S.L. (2019) Graph-based genome alignment and genotyping with HISAT2 and HISAT-genotype. Nature Biotechnology, 37, 907–915.

18. Kim, D., Pertea, G., Trapnell, C., Pimentel, H., Kelley, R., and Salzberg, S.L. (2013) TopHat2: accurate alignment of transcriptomes in the presence of insertions, deletions and gene fusions. Genome Biology, 14, R36.

19. Kozieł, E., Otulak-Kozieł, K., and Rusin, P. (2024) Glutathione-the “master” antioxidant in the regulation of resistant and susceptible host-plant virus-interaction. Frontiers in Plant Science, 15, 1373801.

20. Langmead, B. and Salzberg, S.L. (2012) Fast gapped-read alignment with Bowtie 2. Nature Methods, 9, 357–359.

21. Li, S., Yamada, M., Han, X., Ohler, U., and Benfey, P.N. (2016) *High-resolution expression map of the* Arabidopsis *root reveals alternative splicing and lincRNA regulation*. Developmental Cell, 39, 508–522.

22. Liu, G.-T., Wang, B.-B., Lecourieux, D., Li, M.-J., Liu, M.-B., Liu, R.-Q., et al. (2021) *Proteomic analysis of early-stage incompatible and compatible interactions between grapevine and* P. viticola. Horticulture Research, 8, 100.

23. Love, M.I., Huber, W., and Anders, S. (2014) Moderated estimation of fold change and dispersion for RNA-seq data with DESeq2. Genome Biology, 15, 550.

24. Mohr, P.G. and Cahill, D.M. (2007) *Suppression by ABA of salicylic acid and lignin accumulation and the expression of multiple genes, in* Arabidopsis *infected with* Pseudomonas syringae pv. tomato. Functional & Integrative Genomics, 7, 181–191.

25. Musich, R., Cadle-Davidson, L., and Osier, M.V. (2021) Comparison of short-read sequence aligners indicates strengths and weaknesses for biologists to consider. Frontiers in Plant Science, 12, 657240.

26. Noctor, G. and Foyer, C.H. (1998) ASCORBATE AND GLUTATHIONE: keeping active oxygen under control. Annual Review of Plant Physiology and Plant Molecular Biology, 49, 249–279.

27. Panchy, N., Lehti-Shiu, M., and Shiu, S.-H. (2016) Evolution of gene duplication in plants. Plant Physiology, 171, 2294–2316.

28. Putri, G.H., Anders, S., Pyl, P.T., Pimanda, J.E., and Zanini, F. (2022) Analysing high-throughput sequencing data in Python with HTSeq 2.0. Bioinformatics, 38, 2943–2945.

29. Raplee, I.D., Evsikov, A.V., and Marín De Evsikova, C. (2019) Aligning the Aligners: comparison of RNA sequencing data alignment and gene expression quantification tools for clinical breast cancer research. Journal of Personalized Medicine, 9, 18.

30. Reineke, G., Heinze, B., Schirawski, J., Buettner, H., Kahmann, R., and Basse, C.W. (2008) *Indole-3-acetic acid (IAA) biosynthesis in the smut fungus* Ustilago maydis *and its relevance for increased IAA levels in infected tissue and host tumour formation*. Molecular Plant Pathology, 9, 339–355.

31. Robinson, J.T., Thorvaldsdottir, H., Turner, D., and Mesirov, J.P. (2023) igv.js: an embeddable JavaScript implementation of the Integrative Genomics Viewer (IGV). Bioinformatics, 39, btac830.

32. Ruan, X., Ma, L., Zhang, Y., Wang, Q., and Gao, X. (2021) *Dissection of the complex transcription and metabolism regulation networks associated with maize resistance to* Ustilago maydis. Genes, 12, 1789.

33. Ryu, K.H., Huang, L., Kang, H.M., and Schiefelbein, J. (2019) Single-cell RNA sequencing resolves molecular relationships among individual plant cells. Plant Physiology, 179, 1444–1456.

34. Shimizu, T. and Suzaki, K. (2020) Multiple gene clusters responsive to Plasmopara viticola infection in grapevines. European Journal of Plant Pathology, 158, 681–691.

35. Song, H., Niu, T., Dong, J., Hou, Q., Feng, Y., Liu, H., et al. (2025) Tandem duplicate genes preferentially contribute to the resistance of abiotic stress in sunflowers. Industrial Crops and Products, 230, 121129.

36. Stark, R., Grzelak, M., and Hadfield, J. (2019) RNA sequencing: the teenage years. Nature Reviews Genetics, 20, 631–656.

37. Trapnell, C., Pachter, L., and Salzberg, S.L. (2009) *TopHat: discovering splice junctions with RNA-Seq*. *Bioinformatics (Oxford*, England), 25, 1105–1111.

38. Wang, P., Moore, B.M., Panchy, N.L., Meng, F., Lehti-Shiu, M.D., and Shiu, S.-H. (2018) Factors Influencing Gene Family Size Variation Among Related Species in a Plant Family, Solanaceae. Genome Biology and Evolution, 10, 2596–2613.

39. Wang, Y., Li, J., and Paterson, A.H. (2013) MCScanX-transposed *: detecting transposed gene duplications based on multiple colinearity scans*. Bioinformatics, 29, 1458–1460.

40. Wang, Z., Gerstein, M., and Snyder, M. (2009) RNA-Seq: a revolutionary tool for transcriptomics. Nature Reviews Genetics, 10, 57–63.

41. Xu, S., Hu, E., Cai, Y., Xie, Z., Luo, X., Zhan, L., et al. (2024) Using clusterProfiler to characterize multiomics data. Nature Protocols, 19, 3292–3320.

42. Yang, Y., Li, Q., Li, H., Qi, K., Xie, Z., Wang, Z., et al. (2025) Comprehensive identification and analysis of clusters of tandemly duplicated genes reveal their contributions to adaptive evolution of green plants. The Plant Journal, 123, e70370.

